# Production of a functionally active recombinant SARS-CoV-2 (COVID-19) 3C-Like protease and a soluble inactive 3C-like protease-RBD chimeric in a prokaryotic expression system

**DOI:** 10.1101/2022.03.25.485815

**Authors:** Carolina De Marco Verissimo, Jesus Lopez-Corrales, Amber L. Dorey, Krystyna Cwiklinski, Richard Lalor, Nichola Eliza Davies Calvani, Heather L. Jewhurst, Sean Doyle, John P. Dalton

## Abstract

During the SARS-CoV-2 intracellular life-cycle, two large polyproteins, pp1a and pp1ab, are produced. Processing of these by viral cysteine proteases, the papain-like protease (PLpro) and the chymotrypsin-like 3C-like protease (3CL-pro) release non-structural proteins necessary for the establishment of the viral replication and transcription complex (RTC), crucial for viral replication. Hence, these proteases are considered prime targets against which anti-COVID-19 drugs could be developed. Here, we describe the expression of a highly soluble and functionally active recombinant 3CL-pro using *Escherichia coli* BL21 cells. In addition, we assessed the ability of our 3CL-pro to function as a carrier for the Receptor Binding Domain (RBD) of the Spike protein. The co-expressed chimeric protein, 3CLpro-RBD, did not exhibit 3CL-pro activity, but its enhanced solubility made purification easier and improved RBD antigenicity when tested against serum from vaccinated individuals in ELISAs. When used to immunise mice, the 3CLpro-RBD chimer elicited antibodies mainly to the 3CL-pro portion of the molecule indicating that a different chimeric composition (i.e., RBD/full Spike-3CLpro) or expression system (i.e., mammalian cells), might be required to produce and deliver a RBD with immunogenicity similar to the native protein. Chimeric proteins containing the 3CL-pro could represent an innovative approach to developing new COVID-19 vaccines.

## INTRODUCTION

The severe acute respiratory syndrome coronavirus (SARS-CoV-2) emerged in Wuhan, China, in December 2019 and quickly spread throughout the world [1–3]. Person-to-person transmission of the virus resulted in rapid distribution of SARS-CoV-2, leading to the unprecedented pandemic of coronavirus disease 2019 (COVID-19), which up to now has claimed >6 million lives [4]. The impact of this pandemic on the global health and economy prompted the rapid action on the development, testing and approval of prophylactic COVID-19 vaccines, followed by mass immunization programs [5].

SARS-CoV-2 is an enveloped virus that contains a single-strand of positive-sense RNA. Infection begins when the virus attaches to cells via the angiotensin converting enzyme 2 (ACE2) receptor, mediated by the receptor binding domain (RBD) of the major glycoprotein expressed on the virus surface, the Spike protein [6, 7]. Fusion of the viral membrane with the lumen of the endosomal membrane leads to endocytosis, facilitating infection via entry of the viral RNA into the cytosol. Applying new approaches and technologies, a multitude of vaccines have been developed, four of which were licensed by the regulatory agencies [4] and have been administrated across the world, representing a relief and a unique opportunity to prevent the deaths of millions of people and control the pandemic. In general, each of these four vaccines induce antibodies against the Spike protein and bind to the RBD to block its interaction with ACE2 [8, 9].

During the intracellular viral life cycle, two large polyproteins, pp1a and pp1ab, are translated. Sixteen non-structural proteins (nsp) are co-translationally and post-translationally released from pp1a and pp1ab upon proteolytic activity of two virus cysteine proteases, the papain-like protease (PLpro) and the chymotrypsin-like 3C-like protease (3CL-pro), also known as the main protease. These proteases allow the establishment of the viral replication and transcription complex (RTC), which is crucial for virus replication inside the cells [10]. The 3CL-pro plays a prominent role on viral gene expression and replication [11, 12]. Moreover, recent studies comparing the protease from other coronaviruses, SARS-CoV and Middle East Respiratory Syndrome CoV (MERS-CoV), or from picornaviruses, show that they are all highly conserved in terms of proteolytic activity and structure [13–15] and revealed important immunomodulatory properties for this enzyme. Amongst other mechanisms, during viral infection 3CL-pro contributes to the delay of host anti-viral innate immune response by cleaving or inactivating key elements of the Retinoic acid-inducible gene I (RIG-I) like receptors (RLRs)-mediated Type I interferon (INF-I) signalling pathway, which allows effective viral infection and contribute for disease progression and severity [15–17]. Thus, the 3CL-pro could be considered an attractive target for the development of future anti-COVID-19 treatments.

Here we describe the production of a recombinant 3CL-pro in a prokaryotic expression system and its purification as a highly soluble and functionally active protease. We also generated a 3C-like protease-RBD gene construct that enabled the production of a chimeric protein, named 3CLpro-RBD. This strategy proved useful to enhance the solubility and antigenicity of the RBD, albeit the recombinant chimeric protein did not exhibit proteolytic activity (understandable since the functional 3CL-pro functions as a dimer). When used to immunise mice, the chimeric protein elicited antibodies mainly to 3CL-pro indicating that alternative chimeric format or expression system might be required to produce an antigenic RBD. Nonetheless, the use of SARS-CoV-2 chimeric proteins containing 3CL-pro could represent an innovative way to develop alternative COVID-19 vaccines that may induce both humoral and cellular immune responses.

## METHODS

### Ethical statement

Human experimental work was conducted according to Human Research Ethics Committees. Sera samples from individuals double-vaccinated with Pfizer/BioNTech (BNT162b2) vaccine were obtained from healthy volunteers following ethical approval by the National University of Ireland Galway, Ireland, research ethics committee (R20.Jun.06). The samples were pooled and immediately stored at −80°C. All participants provided written informed consent prior to the study. Negative control samples obtained from the Irish Blood Transfusion Service. These blood samples were previously characterized by De Marco Verissimo et al. [18].

### Recombinant protein production in *Escherichia coli* cells and purification

Sequences encoding the 3CL-pro and RBD proteins were codon optimized for expression in *Escherichia coli* and cloned into the pET-28a(+) vector (Genscript Biotech). The chimeric protein 3CLpro-RBD was produced by generating a gene construct that linked the 3CL-pro and RBD genes by a bridge sequence that encoded for glycine-proline triple repeat (GPGPGP) (see Figure 1). The recombinantly produced proteins contain a thrombin cleavage site followed by a C-terminal His-tag. The synthesized vectors were transformed into BL21 competent *E. coli* cells (ThermoFisher Scientific) following the manufacturer’s instructions and stored in Luria Bertani (LB) broth (Sigma-Aldrich) supplemented with 25% glycerol at −80°C. LB broth supplemented with 50 μg/mL kanamycin was inoculated from the glycerol stock and incubated shaking (200 rpm) at 37°C overnight. The culture was then diluted in fresh LB broth supplemented with kanamycin, incubated at 37°C to OD_600_ 0.6 and protein expression induced with 1 mM isopropyl-β-D-1-thiogalactopyranoside (IPTG; ThermoFisher Scientific) for 4 hr at 30°C (3CL-pro and RBD); 18 h at 16°C (3CLpro-RBD chimer). Following centrifugation at 10,000 x *g* for 10 min at 4°C, the bacterial pellets were re-suspended in 10 mL ST buffer (10 mM Tris, 150 mM NaCl, pH 8.0).

**Figure 1.**
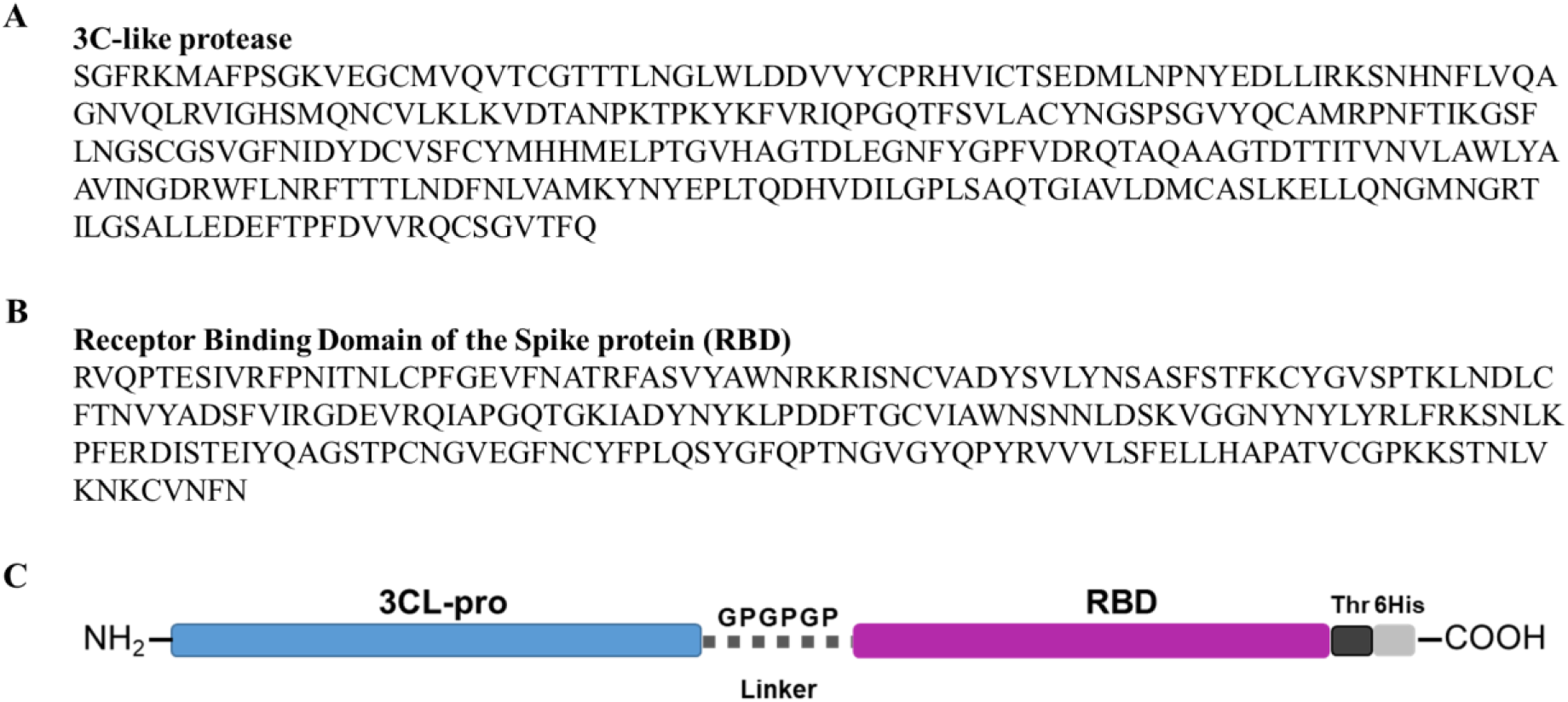
Primary sequence of the SARS-CoV-2 proteins and schematic representation of the 3CLpro-RBD chimeric protein structure. A: The amino acid sequence of the SARS-CoV-2 3C-like protease (3CL-pro) used for recombinant expression in *Escherichia coli.* B: The amino acid sequence of the receptor binding domain (RBD) (residues 319-542 of the full SARS-CoV-2 Spike protein). C: Schematic representation of the 3CLpro-RBD chimeric protein structure showing the unique GP linker. SARS-CoV-2 proteins, 3CL-pro and the RBD are linked by a GP triplet (Glycine, G, and Proline, P), allowing their expression as a stable chimeric protein. Thr: Thrombin cleavage site; 6His: Histidine tag added to the protein C-terminal.

The bacteria pellets were treated with lysozyme (10 μg/mL), sonicated on ice (6 x 10 seconds, 40% amplitude) and centrifuged 15,000 x *g* at 4°C for 30 min. The soluble recombinant protein within the supernatant was purified and dialysed using the Profinia Affinity Chromatography Protein Purification System (Bio-Rad), with the mini profinity IMAC and mini Bio-Gel P-6 desalting cartridges (Bio-Rad). The protein concentration and purity were verified by Bradford Protein Assay (Bio-Rad) and by 4-20% SDS-PAGE gels (Bio-Rad) stained with Biosafe Coomassie (Bio-Rad), respectively. The gels were visualised using a G:BOX Chemi XRQ imager (Syngene).

The recombinant 3CL-pro purified by affinity chromatography was additionally purified using size-exclusion (gel filtration) chromatography to resolve its dimerization state. The purification was performed using a high performance Superdex 75 10/300 GL (Tricorn) column, with a flow rate of 400 μL/min and eluted into 1x PBS. Three known proteins of different molecular sizes were resolved in the column as standards, namely conalbumin (75 kDa), carbonic anhydrase (29 kDa) and aprotinin (6.5 kDa) (Sup Figure S1). Once the retention parameters were determined, the r3CL-pro, at 1 mg/mL in PBS, was added to the column for purification. Aliquots of 200 μL of the sample were collected and stored at 4°C for further analysis using an activity assay (see below).

As RBD protein was found within the inclusion bodies, processing of the pellets, protein purification and dialysis were performed as described by Schlager et al. [19] and employed by us previously to extract recombinant SARS-CoV-2 proteins previously [18]. Briefly, 1% (*w/v*) SDS buffer (8 mM Na_2_HPO_4_, 286 mM NaCl, 1.4 mM KH_2_PO_4_, 2.6 mM KCl, 1% (*w/v*) SDS, pH 7.4) containing 0.1 mM DTT was added to the cell pellet to solubilize the inclusion bodies. After sonication, the samples were centrifuged 15,000 x *g* at 4°C for 30 min and the resulting supernatant containing the target protein was filtered and purified using a pre-equilibrated Ni-NTA beads column (Qiagen). The recombinant protein was eluted using 4 mL of elution buffer (8 mM Na_2_HPO_4_, 286 mM NaCl, 1.4 mM KH_2_PO_4_ 2.6 mM KCl, 0.1% Sarkosyl (*w/v*), 250 mM imidazole, pH 7.4) and buffer-exchanged into 1x PBS containing 0.05% sarkosyl, pH 7.4.

### Fluorogenic assay to assess the enzymatic activity of the recombinant 3CL-pro

The enzymatic activity of the recombinant r3CL-pro purified by affinity chromatography and of the different fractions produced by gel filtration were verified using a fluorogenic assay using the substrate LGSAVLQ-rhodamine 110-dp (BostonBiochem). Unless highlighted, all the screening assays were performed at 37°C, in a 100 μL reaction volume Hepes buffer (20 mM Hepes, 2 mM EDTA, pH 7.4). Initially, the reaction buffer was mixed with either of the recombinant proteins, r3CL-pro (500 nM), rRBD (500 nM) or r3CLpro-RBD chimer (500 nM), and incubated for 5 min at room temperature. The fluorogenic substrate (20 μM) was added to the wells and the proteolytic activity was measured at 37°C, over 1 h, as relative fluorescent units (RFU) in a PolarStar Omega Spectrophotometer (BMG LabTech). All assays were carried out in triplicate. Commercial broad-spectrum protease inhibitors, namely serine protease inhibitors AEBSF (5 mM; Sigma-Aldrich) and Futhan-175 (FUT-175, 200 μM; BD-Pharmingen-Bioscience), and the cysteine protease inhibitor E-64 (200 μM; Sigma-Aldrich), were added to the reaction, individually, for further characterization of the proteolytic activity of the recombinant r3CL-pro.

### Assessment of the immunogenicity of the r3CL-pro, the rRBD and the chimeric r3CLpro-RBD

Seven weeks-old male and female CD1 outbred mice were used to assess the immunogenicity of the recombinantly-produced proteins according to the schedule shown in Figure 2. All animal experimental procedures were carried out by Eurogentec, BE, as follows: Group 1, adjuvant control group (Montanide ISA 206VG, Seppic) (n= 9); Group 2, r3CLpro-RBD chimer (15 μg) formulated in the Montanide adjuvant (1: 1 *v*/*v*) (n=10); Group 3, r3CL-pro (15 μg) formulated in the Montanide adjuvant (1:1 *v*/*v*) (n=7); Group 4, rRBD (15 μg) formulated in the Montanide adjuvant (1:1 *v*/*v*) (n=7); Group 5, r3CL-pro + rRBD (7.5 μg of each) formulated in the Montanide adjuvant (1:1 *v*/*v*) (n=7).

**Figure 2.**
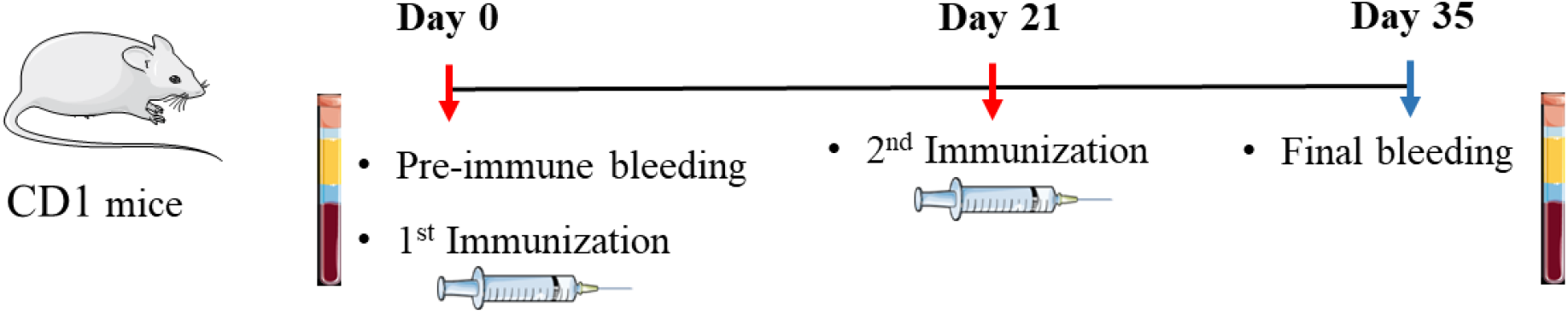
Graphical schematic showing the schedule for the immunization of CD1 outbred mice using the recombinant SARS-CoV-2 proteins. Red arrows indicate when the mice were immunized with adjuvant alone or with the recombinant 3CLpro-RBD chimer, 3CL-pro, RBD, or a mix of 3CL-pro and RBD. Blue arrow indicates the end of the experiment and when the final bleeding was taken.

### ELISA to assess antibodies against the recombinant SARS-CoV-2 proteins in serum of vaccinated humans

Flat-bottom 96 well microtitre plates (Nunc MaxiSorp, Biolegend) were coated with r3CL-pro, r3CLpro-RBD chimer, rRBD or cmRBD as described above. After incubation in blocking buffer (PBST) and washing steps, pooled serum samples from (a) 10 vaccinated individuals (collected at least 10 days after the second dose Pfizer/BioNTech (BNT162b2) vaccine), or from 10 negative controls individuals (samples from the Irish Blood Transfusion Service obtained before COVID-19 pandemic) were diluted 1:100 in blocking buffer and added to the plate. After 1 h incubation at RT, and washing five times with PBST, the secondary antibody HRP anti-Human IgG (Fc specific) (Sigma-Aldrich) was added (1:15,000), and the plates incubated for 1 h at RT. After washing five times, TMB (3,3’,5,5’-Tetramethylbenzidine Liquid Substrate Supersensitive, Sigma-Aldrich) substrate was added to each well. Following a three-minute incubation the reaction was stopped with 2 N sulphuric acid and plates read at 450 nm in a PolarStar Omega Spectrophotometer. All samples were analysed in triplicate.

### Analysis of the immune response of mice to the recombinant SARS-CoV-2 proteins by ELISA and immunoblots

The antibody response of individual mouse serum at day 0 and day 35 was assessed by ELISA, using the following recombinant antigens: r3CLpro-RBD chimer; r3CL-pro; rRBD; recombinant Subunit 2 of the Spike protein (rS2Frag; produced in De Marco Verissimo et al. [18]); commercial RBD produced in mammalian cells (cmRBD, Sigma-Aldrich).

Flat-bottom 96 well microtitre plates (Nunc MaxiSorp, Biolegend) were coated overnight at 4°C with either r3CLpro-RBD chimer (2 μg/mL); r3CL-pro (2 μg/mL); rRBD (2 μg/mL); rS2Frag (1 μg/mL); cmRBD (1 μg/mL) diluted in carbonate buffer (pH 9.6). After incubation in blocking buffer (2% BSA in PBS-0.05% Tween-20 (*v/v*), pH 7.4; PBST) and washing steps, mice serum diluted 1:100 in blocking buffer was added to the antigen-coated wells and incubated for 1 hr at RT. After washing five times with PBST, the secondary antibody HRP goat to mouse-anti-IgG (ThermoFisher Scientific) was added (1:10,000), and the plates incubated for 1 h at RT. After washing five times, TMB substrate was added to each well. Following a three-minute incubation the reaction was stopped with 2 N sulphuric acid and plates read at 450 nm in a PolarStar Omega Spectrophotometer. All samples were analysed in triplicates.

### Detection of neutralizing antibodies in mice sera

The commercially available SARS-CoV-2 Neutralization Antibody Detection Kit (cPass, GenScript) was used to identify the presence of RBD neutralizing antibodies in the serum samples from mice immunized with r3CL-pro, r3CLpro-RBD chimer, rRBD, r3CL-pro+rRBD or Adjuvant. Initially, pooled serum samples from animals in the same group (Day 35) were used for screening. Further analysis of individual samples from mice immunized with r3CLpro-RBD chimer and with rRBD was performed. Samples and controls were analysed in duplicate, following the manufacturer’s instructions.

## RESULTS

### Production of SARS-CoV-2 recombinant proteins in *E. coli*

The 3C-like protease (r3CL-pro) was readily produced as a recombinant protein in *E. coli*; analysis of bacterial lysate showed that it was a prominent protein that separated into the soluble fraction making it easy to isolate by affinity chromatography. The purified protein resolved at the expected molecular size of ~34 kDa, as a highly soluble protein, and our purification yielded 5.3 mg enzyme per litre of bacterial culture (Figure 3A).

**Figure 3.**
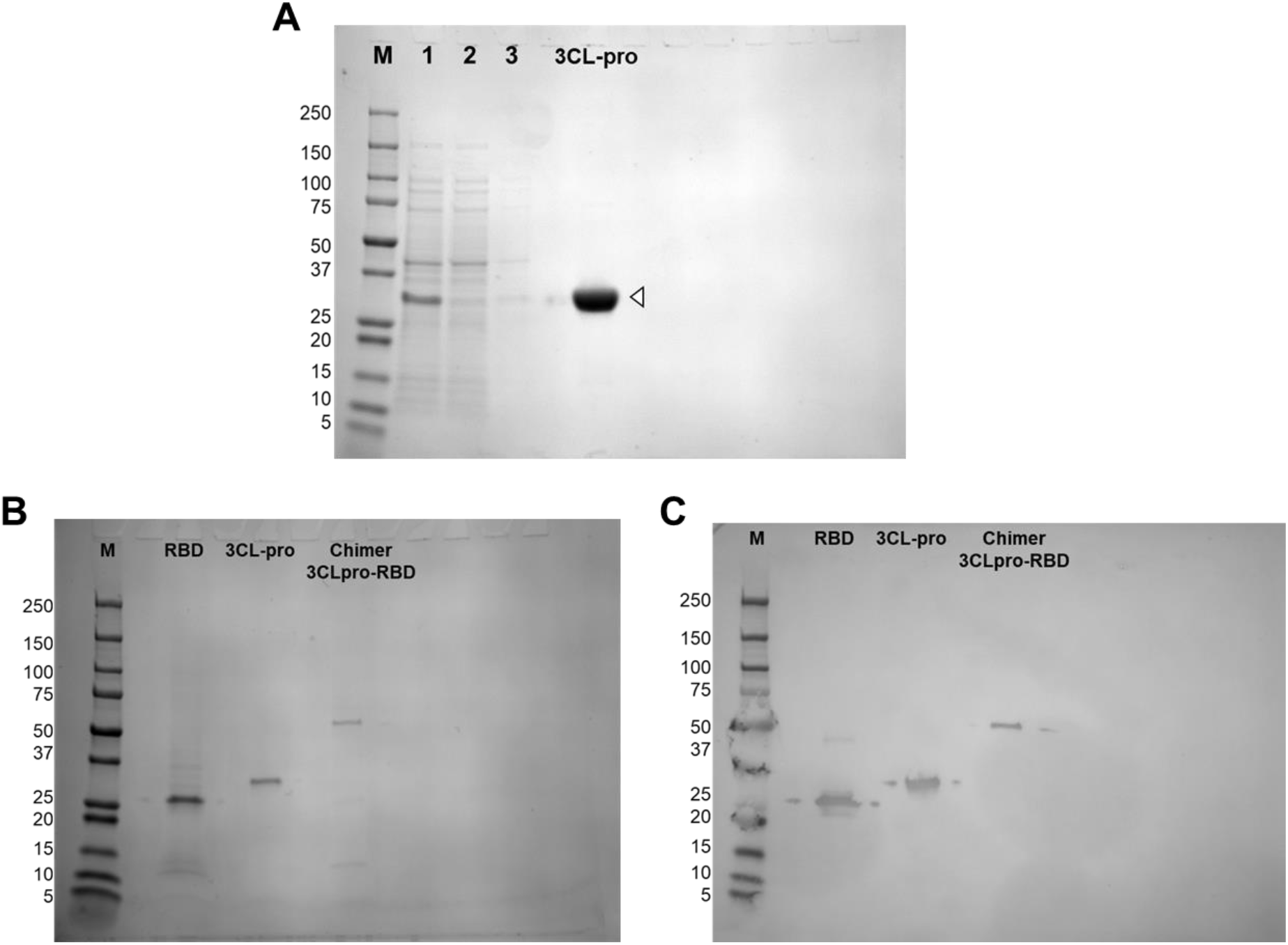
Recombinant expression of the SARS-CoV-2 proteins, 3C-like protease, receptor biding domain (RBD), and 3CLpro-RBD chimer. A: Purification of recombinant 3C-like protease. The supernatant after bacterial pellet digestion (1); proteins that did not bind to the column in the run through (2); proteins in the wash (3); purified and dialysed recombinant protein (3CL-pro). B: The proteins were recombinantly expressed in the prokaryotic expression system, *E. coli,* purified and resolved in SDS-PAGE at the expected respective molecular size: RBD, ~29 kDa; 3CL-pro, ~34 kDa; 3Cpro-RBD chimer, ~60 kDa. B: Western blot of the recombinant proteins probed with the monoclonal anti-6Histidine tag antibody. M: Molecular weight in kilodaltons.

By marked contrast, we found that rRBD did not extract with the solubilisation buffers used but remained in the insoluble pellet, presumably in inclusion bodies. Accordingly, we employed an alternative means of solubilisation that included the chaotropic detergent sodium dodecyl sulphate (SDS) in the buffer, which proved successful in extracting the protein from the pellet [18, 19]. After this extraction procedure, the recombinant RBD could be isolated by NTA-affinity chromatography (Figure 3B). The purified ~29 kDa protein remained soluble after dialysis against PBS containing 0,05% sarkosyl to remove the SDS detergent. This yielded ~1.5 mg of protein per litre of bacterial culture.

By expressing the 3CL-pro and RBD proteins as a chimer, 3CLpro-RBD (~60 kDa), we found that the recombinant protein exhibits intermediate solubility to the protein expressed alone and, therefore, we were able to purify the chimer using the same automated protocol adopted with the r3CL-pro. It provided a yield of ~1.2 mg per litre of bacterial culture (Figure 3B).

Further confirmation that we had purified the targeted proteins was obtained by western blot analysis, where we probed the purified proteins with antibody to the His-tag present on all three recombinant proteins (Figure 3C).

### Recombinant 3CL-pro of SARS-CoV-2 is only functionally active as a dimer, while the chimeric protein does not exhibit activity

Once we successfully produced a soluble recombinant 3CL-pro, we proceeded to check its proteolytic activity using an appropriate substrate. Since the cleavage site of 3CL-pro is highly unique the commercially available LGSAVLQ-rhodamine 110-dp substrate could be utilized to specifically assay the activity of the recombinant enzyme. The assay revealed that our recombinant enzyme is a functionally active protease at 37°C in neutral pH (Figure 4). Unexpectedly, however, despite 3CL-pro being described as a cysteine protease, we found that the enzyme was not inhibited by the cysteine proteases inhibitors such as E-64, but was susceptible to two broad-spectrum serine protease inhibitors, AEBSF and Futhan-175; the enzyme was completely inhibited by these latter compounds at concentrations of 5 mM and 200 μM, respectively (Figure 4).

**Figure 4.**
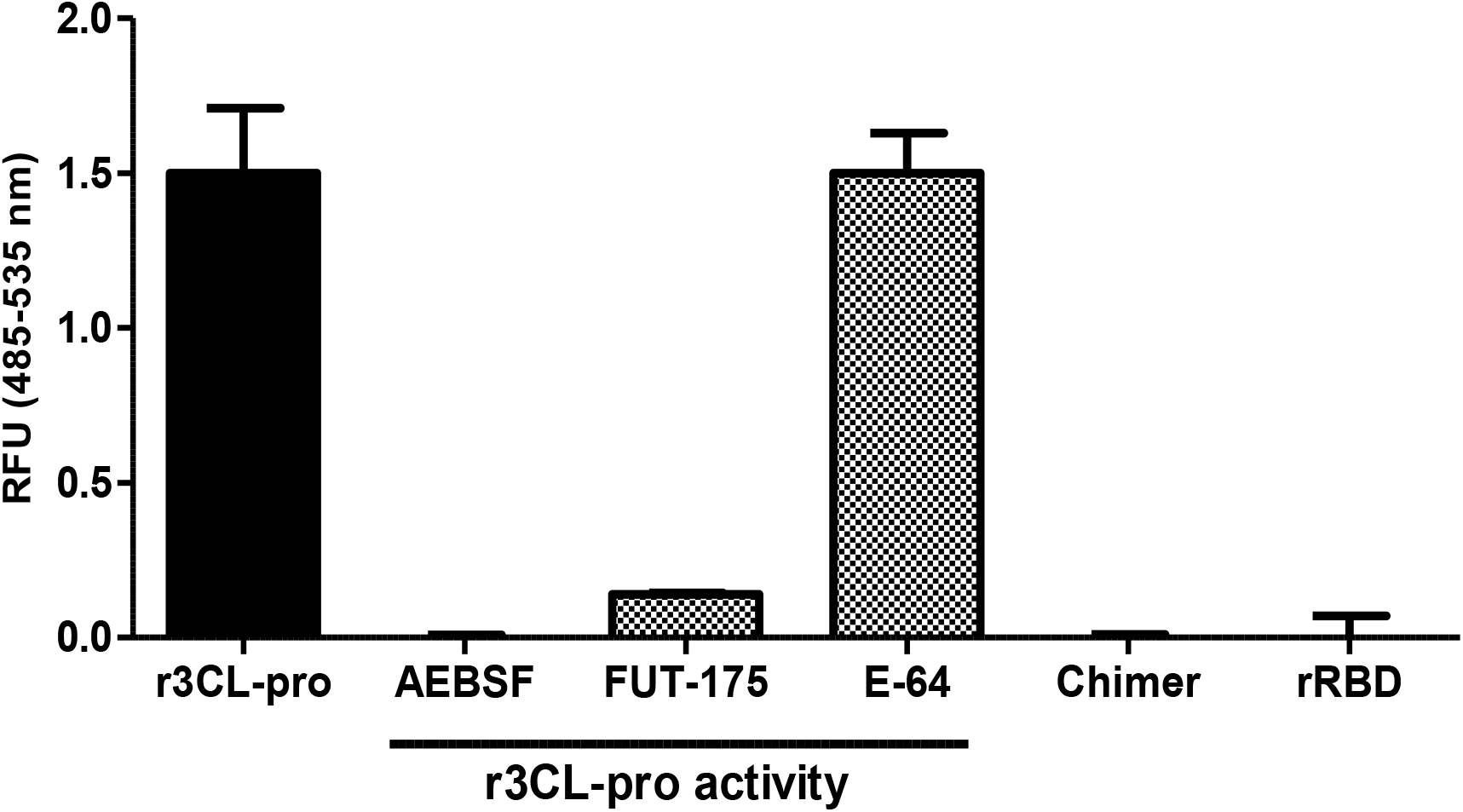
Enzymatic activity of the SARS-CoV-2 recombinant proteins. The enzymatic activity of the r3C-Like protease (r3CL-pro, 500 nM) was tested with or without various broad-spectrum protease inhibitor, namely the serine protease inhibitors AEBSF (5 mM) and Futhan-175 (FUT-175, 200 μM), and the cysteine protease inhibitor E-64 (200 μM). The activity of the r3CLpro-RBD chimer (500 nM) and of the receptor biding domain (rRBD; 500 μM) was assessed in parallel using the same substrate, LGSAVLQ-Rh110 (20 μM). Enzymatic activity presented as relative fluorescence units (RFU) at 485-535 nm. Error bars indicate standard deviation of three separate experiments.

Using gel filtration to further purify the functionally active recombinant r3CL-pro we were able to determine the presence of a mixture of dimers and oligomers within the product purified by affinity chromatography (Peak 1 and 2, respectively; Figure 5). The importance of the dimerization for proteolytic activity of the r3CL-pro was determined by assaying the individual fractions within the two main protein peaks detected during purification. Our data revealed that the r3CL-pro is only functionally active when in its dimeric form, which represents the predominant peak in the chromatogram obtained (Figure 5).

**Figure 5.**
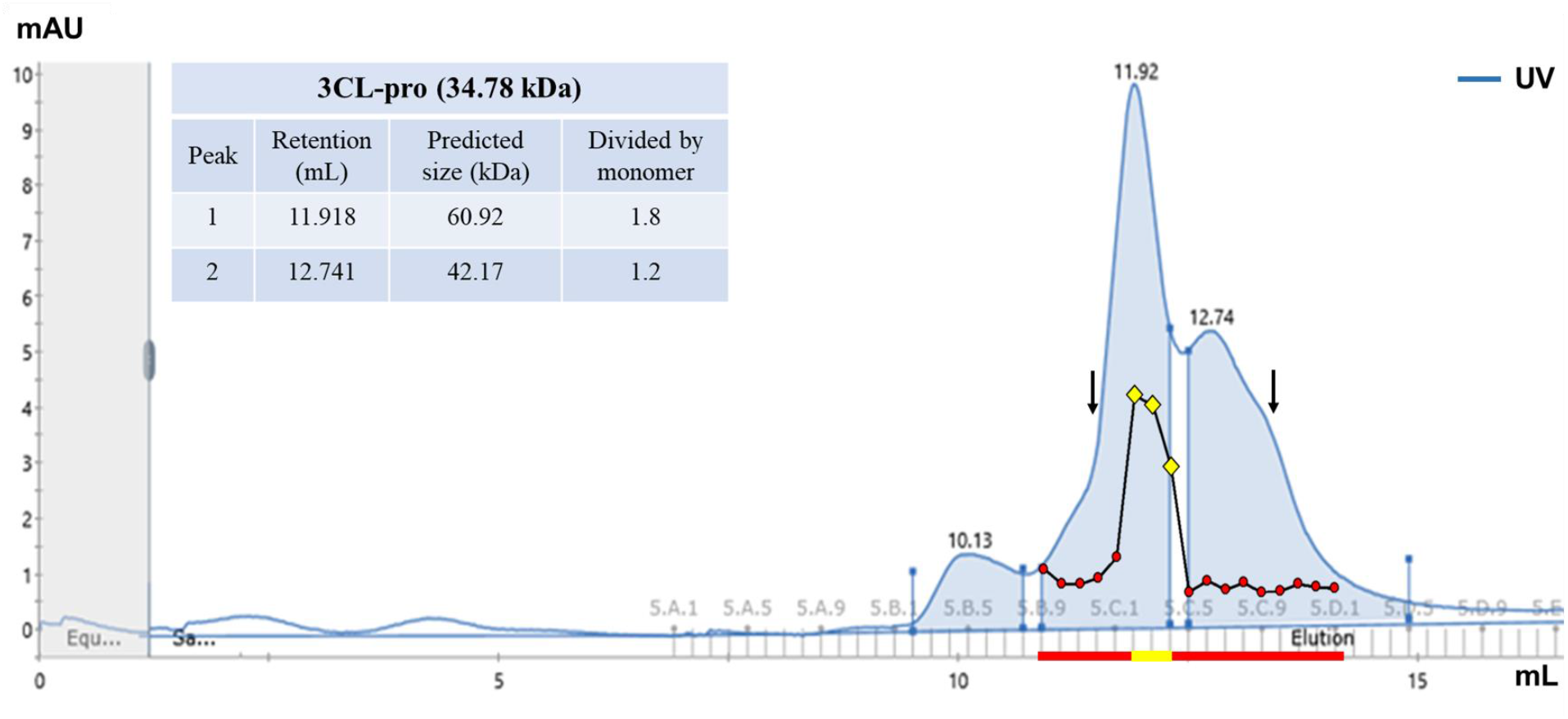
Gel filtration chromatography of the recombinant 3CL-pro. The chromatogram of the purification of the r3CL-pro by gel filtration. Peak 1 (light blue), appeared at 11.918 mL was calculated to represent a protein of ~ 60.9 kDa, while the Peak 2 (light blue), at 12.74 mL, represents a protein of ~ 42 kDa, which indicates the presence of 3CL-pro as a dimer and a oligomer, respectively (for the protein standards data see Supplementary Figure S1). The enzymatic activity of each fraction within the peaks (n = 17, in red) was determined in relation to the activity detected with the r3CL-pro purified only by affinity chromatography. In yellow, the three fractions where enzymatic activity was detected. Black arrows indicate the retention (mL) for the standards conalbumin (11.37) and carbonic anhydrase (13.50). The complete chromatogram for the standards is presented in the Supplementary Fig S2.

In order to examine if the expression of the 3CL-pro in a chimeric format with the RBD, 3CLpro-RBD chimer, had an effect on its proteolytic activity, we also assayed the activity of the recombinant chimeric protein and rRBD produced. Neither protein exhibited enzymatic activity when assayed in the same conditions used with the r3CL-pro (Figure 4).

### Antibodies in serum of naturally infected and vaccinated individuals recognize r3CLpro-RBD chimeric protein

In order to determine if our recombinant SARS-CoV-2 proteins had common epitopes with those present in the virus itself or with the viral proteins expressed upon vaccination, we performed ELISA tests with sera from vaccinated humans using our recombinant proteins r3CL-pro, r3CLpro-RBD chimer and rRBD as target antigens. In parallel, we used the commercial RBD recombinantly produced in mammalian cells (cmRBD), which is commonly used in immunological and functional assays [20]. Our results show that, when compared to negative control samples, sera from vaccinated individuals contain antibodies that recognise the r3CLpro-RBD chimer. These individuals also recognized the rRBD, albeit with lower intensity (Figure 6). Surprisingly, a discrete antibody response against the r3CL-pro was also observed with the vaccinated group (Figure 6).

**Figure 6.**
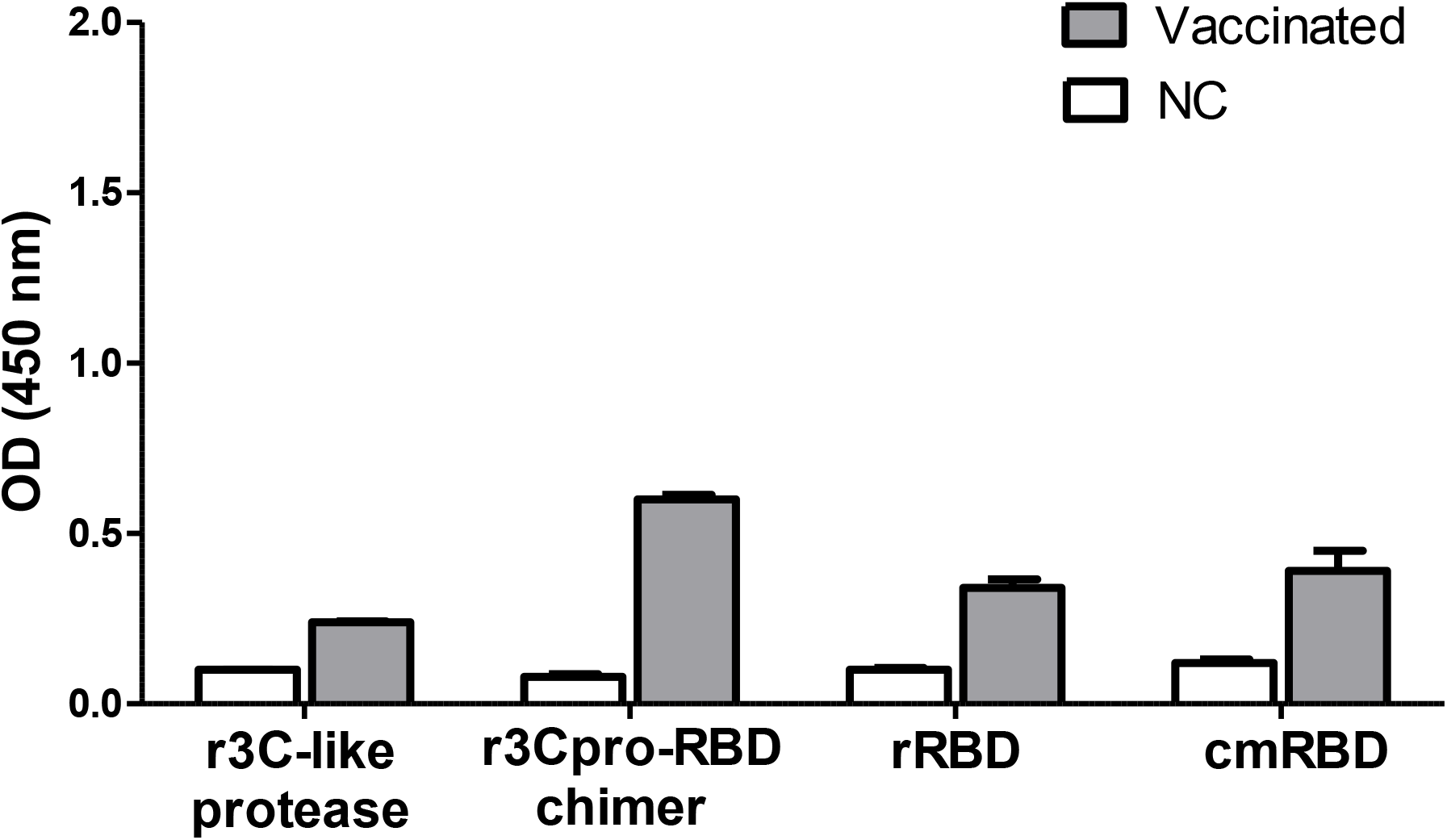
Immune recognition of the recombinant SARS-CoV-2 proteins by antibodies in sera from COVID-19 fully-vaccinated individuals. ELISA tests were performed to assess the presence antibodies in serum of negative control individuals (NC) or COVID-19-vaccinated individuals that bind r3CL-pro, r3CLpro-RBD chimer, rRBD or commercial RBD (cmRBD). Results presented as the mean and standard deviation of OD 450 nm values of all the animals of the group.

### r3CL-pro and r3CLpro-RBD chimer induce antibody response in vaccinated mice

In order to assess and compare the immunogenicity of the recombinant SARS-CoV-2 proteins, we immunized outbred CD1 mice with each protein in a regime similar to that initially recommended for the available COVID-19 vaccines (i.e., initially the recommendation for Pfizer and Oxford vaccines was 2 doses administered 3 weeks a part) [8, 9]. Groups of CD1 mice were immunized with each of the proteins (15 μg) or with a mixture of r3CL-pro and rRBD (7.5 μg each). Finally, as base-line controls, a separated group was vaccinated with adjuvant Montanide ISA 206VG alone, which was used to formulate all the vaccines used in this study. Pre-immune (Day 0) and immune sera (Day 35) of each animal was assessed for antibodies using ELISA tests.

Our ELISA results show that those animals immunized with adjuvant alone did not react to any recombinant protein (Figure 7). They also show that mice immunized with recombinant protein do not react with a non-specific protein, in this case a recombinant fragment of the Spike protein that does not contain the RBD (S2Frag, Figure 7E). Mice immunized with r3CL-pro alone or mixed with rRBD responded with high levels of antibodies against the r3CL-pro but their antibodies did not bind well to the r3CLpro-RBD chimeric (Figure 7A and B). Conversely, mice immunized with the r3CLpro-RBD chimeric produced with antibodies that bound to the chimeric protein in ELISA, but elicited a low level response to the r3CL-pro and the rRBD (Figure 7B). Surprisingly, mice immunized with rRBD did not respond to this antigen or to the r3Cpro-RBD chimeric (Figure 7C). Lastly, mice immunised with rRBD, r3CLpro+RBD or the chimeric protein did not produce antibodies that react with the commercial RBD in ELISA (Figure 7D).

**Figure 7.**
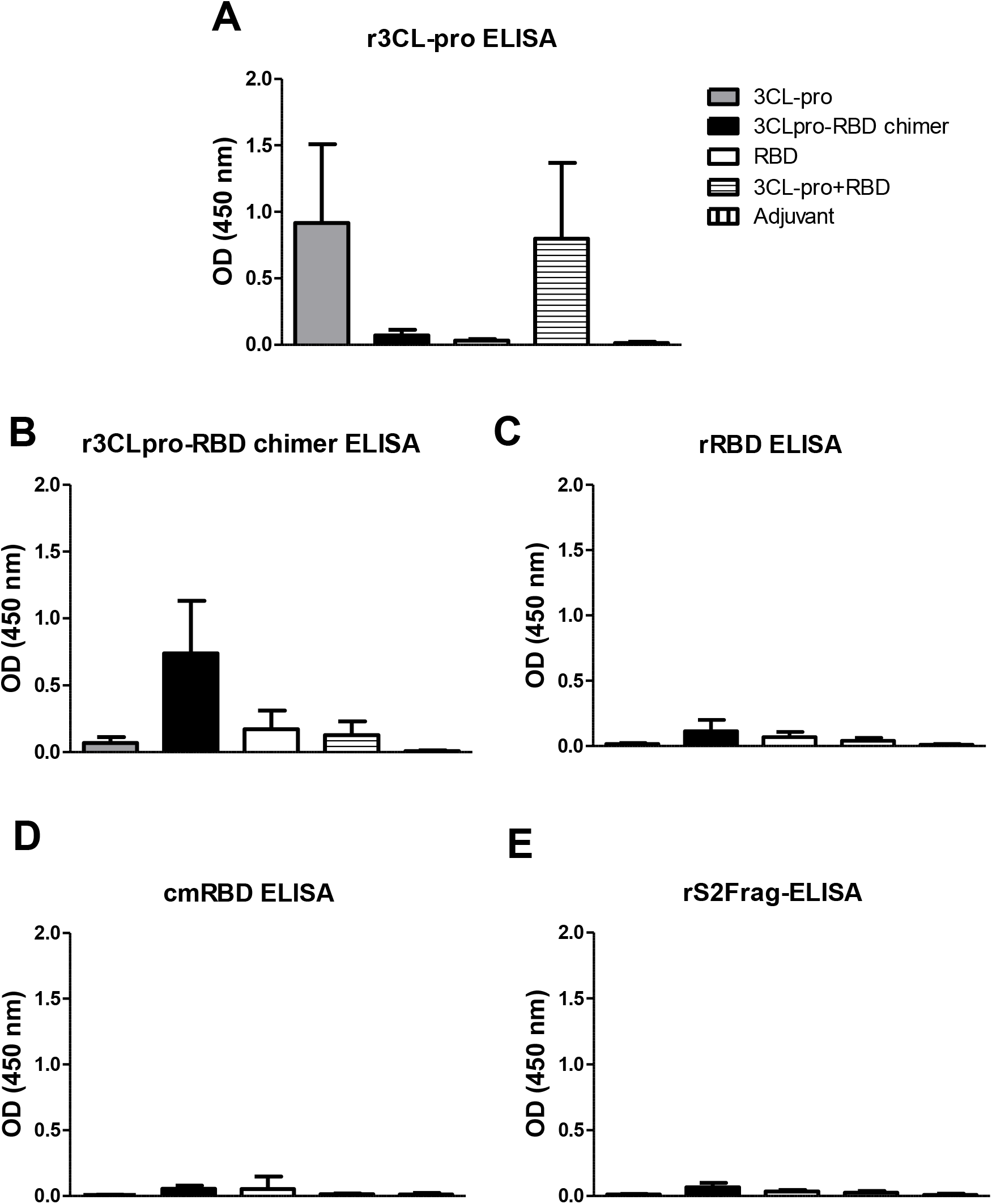
Evaluation of the antibody response induced by the recombinant SARS-CoV-2 proteins in CD1 outbred mice. Groups of CD1 outbred mice were immunized with either r3CL-pro, r3CLpro-RBD chimer, rRBD, r3CL-pro+rRBD or Adjuvant only and evaluated for specific antibodies using ELISA tests. A: ELISA using r3CL-pro as target antigen; B: ELISA using r3CLpro-RBD chimer as target antigen; C: ELISA using rRBD as target antigen; D: ELISA using commercial RBD (cmRBD) as target antigen; E: ELISA using the recombinant subunit 2 of the Spike protein (rS2Frag) as target antigen to assess specificity of the assays. Results presented as the mean and standard deviation of OD 450 nm values of all the animals of the group.

Finally, we assessed the presence of neutralizing antibodies in serum of the mice vaccinated with our recombinant SARS-CoV-2 proteins. The commercial kit used allows for the detection and quantification of antibodies able to bind to recombinant mammalian RBD molecules and consequently block its interaction with the ACE2 receptors present in the ELISA plates (cPass, GenScript). We found that, despite our recombinant proteins were able to induce antibodies when used to immunise animals, those antibodies were unable to bind and neutralize the RBD interaction with ACE2 receptor in the assay used.

## DISCUSSION

The 3C-like protease is regarded as a prime target for therapeutic drug treatment of COVID-19 due to its unique specificity for cleaving peptide bonds that are absent in human proteins [14, 21, 22]. This protease plays a central role during viral replication, being responsible for the cleavage of 11 sites within the polyprotein 1ab, ultimately releasing 13 non-structural proteins involved in SARS-CoV-2 replication inside the host’s cells [23, 24]. Such activity is associated with the ability of this protease to recognize and cleave the unique peptide sequence Leu/Phe/Met-Gln ↓ Gly/Ser/Ala (↓ denotes the cleavage site). We were able to demonstrate that the r3CL-pro produced in the present study using bacterial expression systems has the same requirement for proteolytic activity, including for a glutamine (Gln) at the P1 position, as the recombinant enzyme was able to cleave specifically the LGSAVLQ-Rh110 substrate. Since 3CL-pro is reported to be functionally active as a dimer [25, 26], we further investigated its molecular state using size-exclusion chromatography and determined that the purified product was dominated by r3CL-pro dimers. Together with the results of the activity assays, our data indicate that the production of this enzyme in the *E. coli* system employed here allowed for protein fold and dimerization similar to that of the native form. The unexpected inhibition of this protease by broad-spectrum serine protease inhibitors, but not by cysteine protease inhibitors, does not support the claim that this protease is a cysteine peptidase, but is more in keeping with the reported chymotrypsin-like protease activity of 3CL-pro [10, 27]. Perhaps this observation could be exploited in anti-3CL-pro novel drug design.

In our previous study aimed at improving COVID-19 diagnostics [18], in parallel with the nucleocapsidic (Npro) and S2Frag proteins as target antigens on ELISA tests, we also evaluated the antibody response of individuals naturally infected with SARS-CoV-2 against the r3CL-pro. Our data showed that, although higher antibodies titres are generated against Npro and S2frag, 50% of the individuals analysed mounted a significant antibody response to 3CL-pro, indicating that this protease is naturally immunogenic during COVID-19 infection (Supplementary Fig S2). In the present study, we demonstrated that the recombinant version of this protein is also immunogenic, and induces a strong and specific antibody response in mice. Together, this data indicates the potential application of the 3CL-pro as a vaccine target against COVID-19. Inducing antibodies to the 3CL-pro could provide broader antibody and cellular immunity to individuals and thus induce a stronger protection against SARS-CoV-2 infection [15, 16].

Given the difficulty in producing recombinant soluble SARS-CoV-2 proteins in prokaryotic systems [18, 28, 29], we considered that the highly soluble 3CL-pro could serve as a carrier protein to produce and deliver RBD. Thus, we designed a construct to generate a recombinant chimeric protein, 3CLpro-RBD. The two proteins were linked using a GP-linker that allows separation and flexibility between them, so that each molecule is stable and can function separately. This approach improved the solubility and production of the RBD. However, we observed that the functional activity of the r3CL-pro chimeric was lost, likely due to incorrect protease folding or the inability of the 3CL-pro to form dimers when linked to the RBD.

Surprisingly, our ELISA tests revealed that CD1 mice immunized with the chimeric protein developed antibodies against the 3CL-pro portion of the antigen, but not to the RBD part. Nonetheless, antibodies in serum from individuals fully-vaccinated against COVID-19 recognized the chimeric protein more efficiently than the rRBD alone, indicating that the presentation of the RBD is improved when produced in the chimeric format. Furthermore, the ability of vaccinated individuals to recognise the commercial RBD produced in mammalian cells (cmRBD), but not our rRBD produced in *E. coli* cells, suggests that glycosylated epitopes are important in antibody recognition. This is reinforced by the absence of antibodies capable of preventing RBD-ACE2 binding in serum from mice immunized with our recombinant proteins.

Based on current data, antibodies that bind to correct portions of the RBD can neutralize the SARS-CoV-2 virus by preventing it binding to ACE2 receptors and consequent host cells invasion and infection [22, 30–32]. Therefore, since this is also the mechanism by which the current BioNTech/Pfizer, Moderna, Oxford/Astrazenca and Janssen vaccines perform [8, 9, 33, 34], our chimeric protein could be re-designed in order to improve the molecule antigenicity. This could include: changing the order of the proteins into an arrangement RBD-3CLpro, combining the 3CL-pro with the full-length Spike protein instead of the RBD, or even choosing a mammalian expression system (e.g. HEK cells) to allow glycosylation of the recombinant proteins, which could help generate molecules able to stimulate desirable neutralizing antibodies.

A chimeric protein carrying the RBD certainly could help to circumvent the low immunogenicity problem faced with this molecule, and the combination of RBD with more immunogenic molecules has been demonstrated to be a useful strategy to create alternative vaccine targets to fight COVID-19 [28, 29, 35]. This is the first time that another SARS-CoV-2 molecule, and specifically the 3CL-pro, has been considered as the RBD carrier. Compared to the strategies so far adopted to develop COVID-19 vaccine targets, fusion molecules containing 3CL-pro could have a superior ability to induce long-term neutralizing antibody responses, as well as potent cellular immunity, and also overcome the problems with the SARS-CoV-2 variants that are mainly associated high plasticity of the Spike protein [34, 36, 37].

## CONCLUSIONS

Here we present a straightforward, efficient and cheap method to express and purify a highly soluble and functional 3C-like protease, which is regarded as a main drug target at which to develop therapies against SARS-CoV-2 infection. The enzyme could be useful in the development of high-throughput assays for the screening and isolation of new anti-COVID-19 compounds. In addition, given its solubility and potential for triggering a cellular immune response to fight infection, we introduced the idea of using the 3CL-pro as a carrier to other SARS-CoV-2 proteins, in this case RBD, to improve their expression and delivery as potential vaccines. Co-expressing 3CL-pro and RBD in a chimeric format resulted in loss of the 3CL-pro activity (likely due to the inability of the enzyme domain to dimerise), but enhanced the solubility of the RBD expressed alone, and improved its antigenic properties. When used to immunise mice, the 3CLpro-RBD chimer elicited antibodies mainly to the 3CL-pro portion of the chimer indicating that a different composition (i.e., RBD/full Spike-3CLpro) or expression system (i.e., mammalian cells) might be required to produce and deliver an immunogenic RBD that can elicit viral blocking antibodies. Chimeric proteins containing the 3CL-pro could represent a new approach to engender next generation protein-subunit COVID-19 vaccine candidates.

## DATA AVAILABILITY AND SUPPLEMENTARY FILE

The data that support the findings of this study are all included in the publication.

Supplementary file data archive available on the Cambridge University Press - Cambridge Core website.

