## Supplementary Figure S1 for "Production of a functionally active recombinant SARS-CoV-2 (COVID-19) 3C-Like protease and a soluble inactive 3C-like protease-RBD chimeric in a prokaryotic expression system"

**SUPPLEMENTARY MATERIAL**

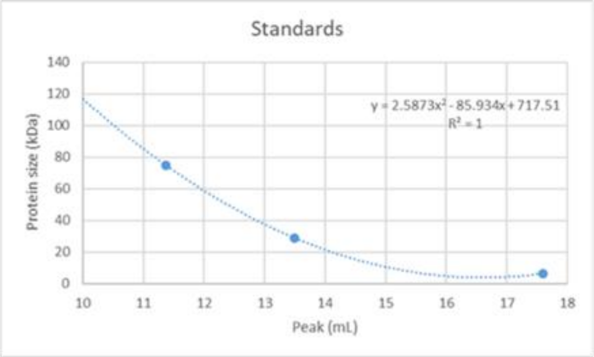


**Peak Table - UV**

| Peak | Retention ml | Area ml*mAU | Area % | Ext coeff. mg⁻¹ ml cm⁻¹ | Fraction(s) | Volume ml | Conductivity mS/cm |
|---|---|---|---|---|---|---|---|
| Peak A | 11.367 | 75.31 | 22.38 | | 5.B.9 - 5.C.5 | 1.801 | 14.81 |
| Peak B | 13.496 | 95.26 | 28.3 | | 5.C.7 - 5.D.5 | 2.200 | 14.80 |
| Peak C | 17.598 | 166.0 | 49.32 | | 5.D.9 - 5.F.4 | 4.001 | 14.79 |

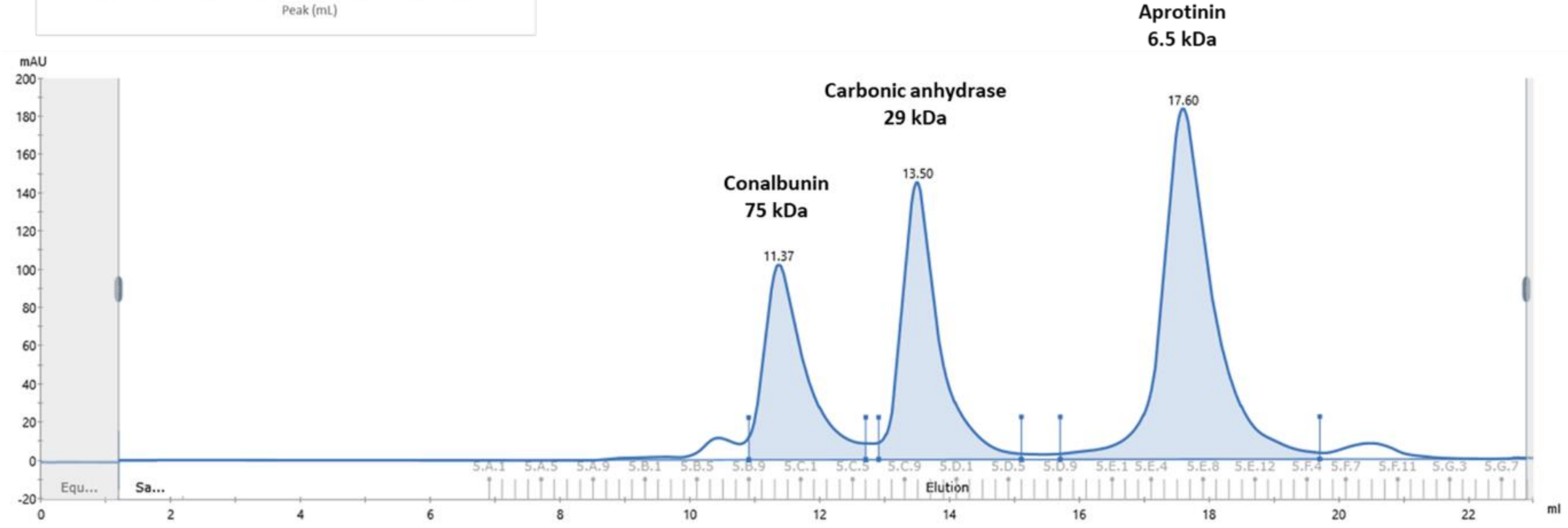


**Supplementary Figure S1.** Protein standards optimization for size exclusion chromatography.
