## Supplementary Figure S2 for "Production of a functionally active recombinant SARS-CoV-2 (COVID-19) 3C-Like protease and a soluble inactive 3C-like protease-RBD chimeric in a prokaryotic expression system"

SUPPLEMENTARY MATERIAL

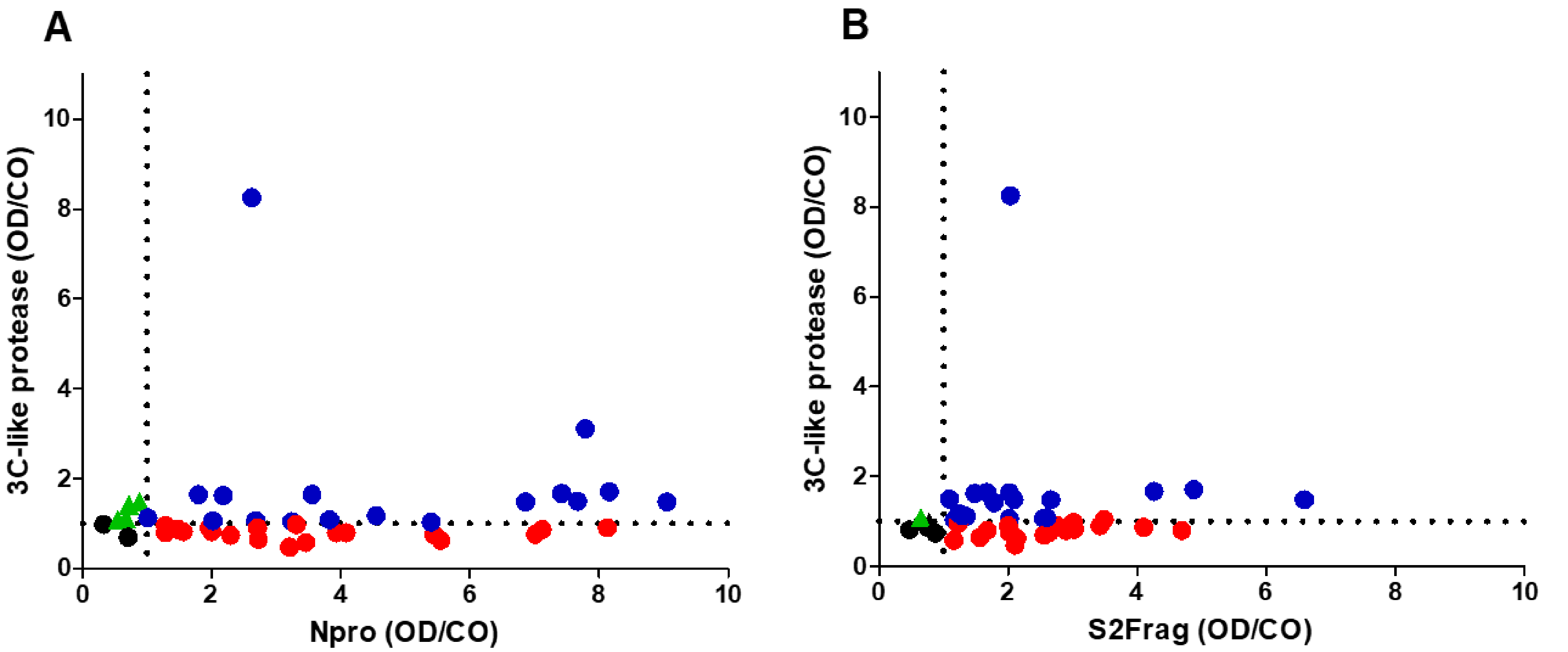


**Supplementary Figure S2. Antibody response of individuals naturally infected with SARS-CoV-2 to the 3C-like protease.** IgG antibodies against the r3C-like protease were detected in sera from 42 individuals confirmed positive for SARS-CoV-2 infection by RT-PCR. (A) Sera from the RT-PCR positive SARS-CoV-2 patients were tested for antibodies against 3C-like protease and compared to their antibodies levels against the Nucleoprotein (Npro). (A) Sera from the RT-PCR positive SARS-CoV-2 patients were tested for antibodies against 3C-like protease and compared to their antibodies levels against the Subunit 2 of the Spike protein (S2Frag). ELISA antibody tests developed in our previous study (De Marco

Verissimo et al., 2021). ▲ sera were negative for antibodies against both Npro and S2frag by ELISA but positive for 3C-like protease; ● sera were negative for antibodies against any of the viral antigens tested; ● sera were positive for antibodies against Npro or S2frag by ELISA and for antibodies against 3C-like protease; ● sera were positive for antibodies against Npro or S2frag by ELISA but not for antibodies against 3C-like protease. Individual ELISA results are presented as Optical density (OD 450 nm) divided by the calculated cut-off (CO) to each ELISA test developed (considering the negative control group). The cut-off value for each antigen is indicated by the dotted line.
